# Autocatalytic assembly of a chimeric aminoacyl-RNA synthetase ribozyme

**DOI:** 10.1101/2024.10.26.620442

**Authors:** Aleksandar Radakovic, Marco Todisco, Anmol Mishra, Jack W. Szostak

## Abstract

Autocatalytic reactions driving the self-assembly of biological polymers are important for the origin of life, yet few experimental examples of such reactions exist. Here we report an autocatalytic assembly pathway that generates a chimeric, amino acid-bridged aminoacyl-RNA synthetase ribozyme. The noncovalent complex of ribozyme fragments initiates low level aminoacylation of one of the fragments, which after loop-closing ligation generates a highly active covalently-linked chimeric ribozyme. The generation of this ribozyme is increasingly efficient over time due to the autocatalytic assembly cycle that sustains the ribozyme over indefinite cycles of serial dilution. Due to its *trans* activity, this ribozyme also assembles ribozymes distinct from itself, such as the hammerhead, suggesting that RNA aminoacylation, coupled with nonenzymatic ligation, could have facilitated the emergence and propagation of ribozymes.

## Introduction

Reactions in which the product catalyzes its own formation are referred to as autocatalytic(*1–3)*. Drawing inspiration from simple, autocatalytic reactions involving small molecules, more complex autocatalytic schemes that produce enzymes that can catalyze their own synthesis, such as an RNA replicase, have been commonly invoked to explain the origin of life(*4–14)*. A long-standing goal of the origin-of-life field has been to identify and harness autocatalytic replicative systems involving genetic and/or catalytic polymers. Indeed, computational simulations suggest that autocatalytic RNA replicators can under certain conditions lead to perpetual propagation of genetic material(*15–17)*. Yet, despite their importance, only a handful of autocatalytic reactions relevant to the evolution of biochemistry have been reported to date.

The earliest examples of biopolymer autocatalysis leveraged modified, palindromic nucleic acids, which generated exact copies of themselves by nonenzymatic templated ligation of two component oligonucleotides(*18–20)*. Autocatalysis in these systems was severely inhibited by the strong binding of the ligated product to the template (i.e. another copy of itself). Relying on ribozyme-mediated templated ligation, a ligase ribozyme was engineered to self-assemble autocatalytically by ligating its own component oligonucleotides(*21*). The inherent product inhibition in this system was initially overcome by minimizing the base-pairing between the template and the substrate. However, true exponential autocatalysis was achieved after engineering the ligase ribozyme to assemble in a cross-catalytic format with two similar but distinct ligase ribozymes replicating each other (*22, 23*). More recently, the *Azoarcus* group I intron was shown to be capable of autocatalytic covalent self-assembly via energy-neutral transesterification reactions. Its self-assembly was driven in the forward direction to ∼20 % full-length product by burying the three key intron recognition base pairs in stem-loops after transesterification(*24*). Although the templating feature is generally associated with nucleic acids, it was also used to drive autocatalytic self-assembly of an alpha-helical peptide that templated a thioester-promoted peptide ligation of its two component peptides(*25–28*). While these templated ligation-based autocatalytic systems demonstrated that biopolymer self-replication is possible, they either cannot or have not been shown to assemble or replicate other useful biopolymers. Other autocatalytic reactions, such as formose reaction networks(*29–31*), self-reproducing giant vesicles(*32*), autocatalytic micelles(*33–35*), and peptide-like replicator fibers(*36–38*), have been invoked to model the origin of biochemical processes. However, their connection to the evolution of extant biochemistry remains unclear.

One of the hallmarks of extant life is the mutual dependence of RNA and proteins in ribosomal translation(*39, 40*), but how this universally conserved machinery emerged remains a mystery(*41, 42*). We recently discovered that loop-closing ligation of aminoacylated RNA can facilitate the assembly of chimeric ribozymes, thereby linking RNA aminoacylation, an inefficient yet key reaction for the origin of translation, to the efficient construction of RNA-based catalysts (**Fig. 1A**)(*43*). Importantly, the loop-closing ligation reaction requires no template, circumventing the formation of catalytically inactive product-template duplexes, which have plagued most RNA-based autocatalytic systems(*19–21, 44, 45*). Therefore, loop-closing assembly of a chimeric aminoacyl-RNA synthetase ribozyme that could aminoacylate its own component RNAs could in principle proceed indefinitely given sufficient substrates (**Fig. 1B**). If this chimeric ribozyme could also aminoacylate other RNA substrates, it could participate not only in self-replication but also in the aminoacylation and assembly of other potentially useful chimeric RNAs.

**Fig. 1.**
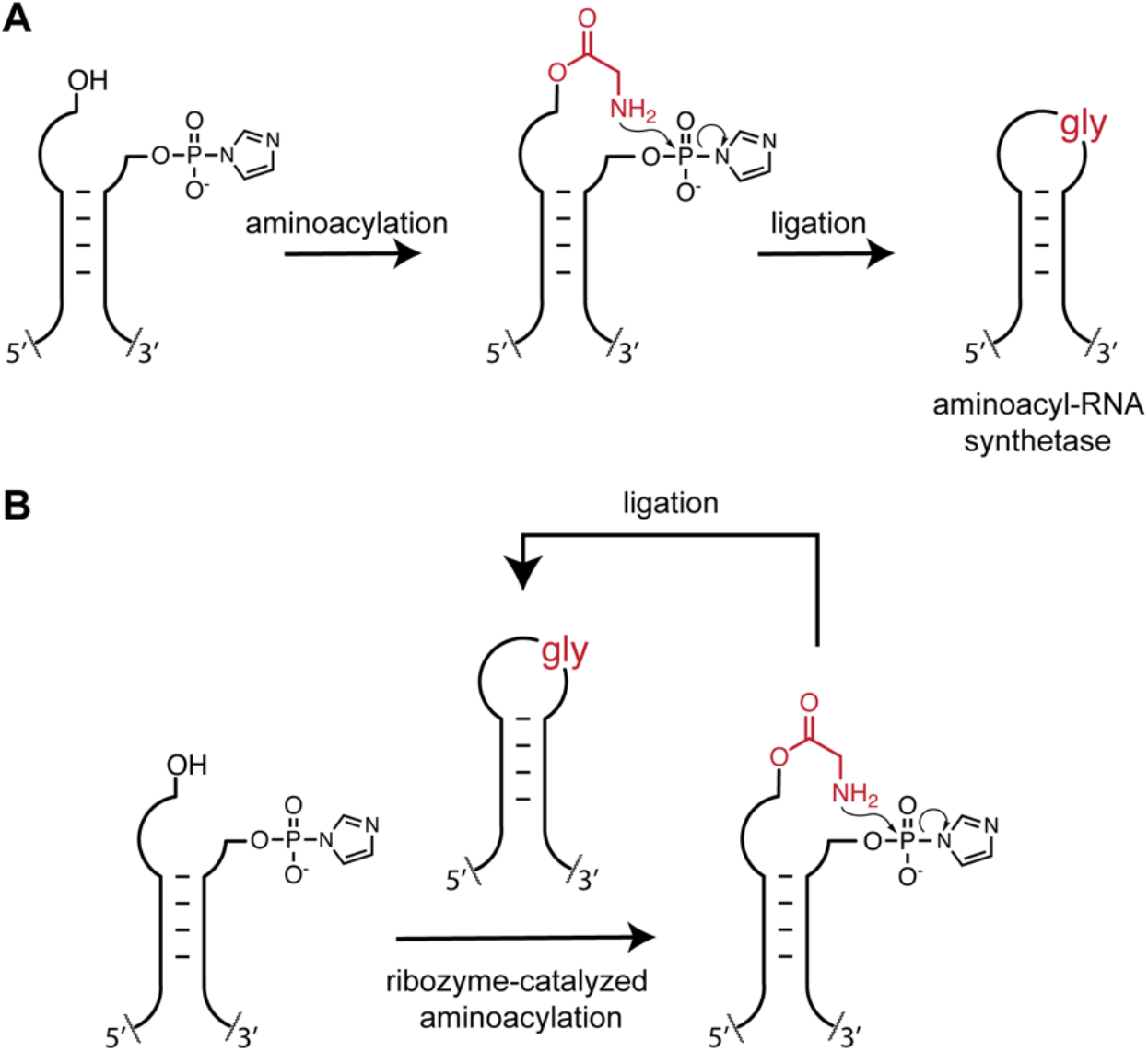
A potential autocatalytic assembly pathway via aminoacylation and loop-closing ligation. **A:** RNA stems with overhangs become aminoacylated either chemically or enzymatically followed by loop-closing ligation mediated by the amino acid to yield covalently linked chimeric hairpin loops. Ligation of appropriate stem-loops leads to a chimeric aminoacyl-RNA synthetase ribozyme (see ref. 43). **B:** The chimeric aminoacyl-RNA synthetase ribozyme catalyzes aminoacylation of the overhang-stems that upon loop-closing ligation make more of the chimeric ribozyme, resulting in autocatalytic self-assembly.

To demonstrate autocatalytic self-assembly, we engineered the Flexizyme aminoacyl-RNA synthetase ribozyme(*46*) to bind and aminoacylate RNA oligonucleotides that upon loop-closure yield another copy of the same chimeric, glycine-bridged Flexizyme. The initial aminoacylation that led to the first copy of the chimeric Flexizyme was performed by the noncovalent assembly of the Flexizyme fragments, which retained sufficient aminoacylation activity to initiate the assembly cycle. We measured the kinetic parameters of each individual reaction in the assembly network and built a global kinetic that accurately predicts the autocatalytic self-assembly cycle. The chimeric Flexizyme was also able to aminoacylate a separate RNA oligonucleotide, which after nonenzymatic loop-closing ligation produced an active, chimeric hammerhead ribozyme, a species distinct from the chimeric Flexizyme. This example shows that a self-replicating RNA-based catalyst can assemble other useful RNA-based catalysts. The evolutionary implication of our work is that aminoacylation of RNA, a reaction normally associated with ribosomal translation, could have been at the center of chimeric ribozyme assembly cycles in the earliest cells. These autocatalytic cycles could have operated as primitive hubs of catalyst assembly within the larger, autocatalytic genome replication cycles, which provided the individual RNA oligonucleotide substrates.

## Results

The Flexizyme contains two stem-loops, P1 and P2, that are distant from the catalytic center, and that can be manipulated with minimal impact on the activity of the ribozyme(*46*). We modified the stem and loop sequences of the dFx Flexizyme(*46*) so that they become substrates for aminoacylation and loop-closing ligation(*43*). We found that the noncovalently assembled fragments of the modified Flexizyme retained a fraction of the aminoacylation activity of the covalent ribozyme (**Figs. S1–2**). Furthermore, the covalent, modified Flexizyme requires only the P1 stem-loop to be ligated to display its full catalytic activity; the P2 stem can remain noncovalently assembled without loss of activity (**Fig. S2**). This result alerted us to the possibility of initiating an autocatalytic assembly cycle by simply mixing the Flexizyme fragments and the Flexizyme substrate, the 3,5-dinitrobenzyl ester of glycine (DBE-gly) (**Fig. S3**). In this proposed cycle, the low-level activity of the noncovalently assembled fragments generates aminoacylated fragments, which after loop-closing ligation result in a highly active covalently linked, chimeric Flexizyme. The chimeric Flexizyme then rapidly aminoacylates more of its component fragments, which ligate to make more of the chimeric Flexizyme, thus driving the autocatalytic assembly cycle (**Figs. 2A and S3**).

**Fig. 2.**
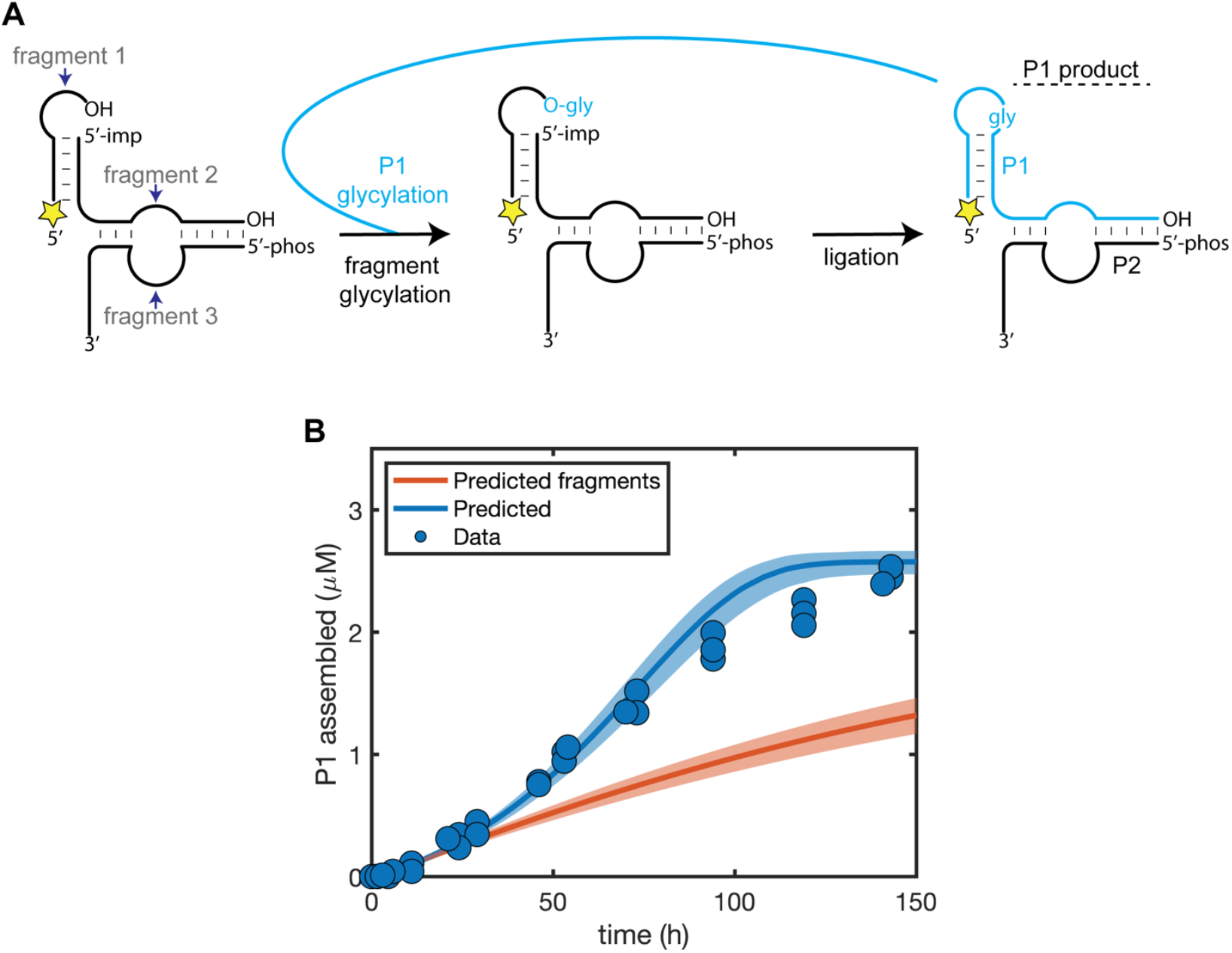
Autocatalytic assembly of the chimeric P1 Flexizyme. **A**: Diagram of the assembly reaction. The fragments self-glycylate via a slow glycylation pathway (black arrow), leading to loop-closing ligation to generate the P1 product. The P1 product then catalyzes a faster glycylation of the fragments (cyan arrow), leading to more P1 (see also **Fig. S3** for sequence details). Yellow star: 5′-FAM label. 5′-imp represents the 5′-phosphorimidazolide. **B**: Assembly of P1 over time with 5 µM of 5′-FAM labeled fragment 1, 5 µM of 5′-phosphorimidazolide fragment 2, and 0.6 µM of fragment 3. The assembly of P1 was monitored by acidic denaturing PAGE in three independent replicates by quantifying the percent of fluorescent, ligated product formed (see **Fig. S4**). The filled circles represent the collected data. The blue trace represents the predicted reaction course from the global kinetic model. The orange trace represents the predicted reaction course if the assembled P1 did not enhance the glycylation and deglycylation reactions. Shaded areas represent 95% confidence intervals for predictions of the model.

By monitoring the assembly of the chimeric P1 Flexizyme from its fragments, we observed a sigmoidal increase in the yield of the P1 ribozyme, consistent with an autocatalytic assembly cycle (**Fig. 2B**, blue circles). To evaluate whether the sigmoidal kinetics were due to autocatalysis and not the multi-step nature of the assembly process, we characterized the kinetics of each reaction in the network independently and built an integrated autocatalytic kinetic model (**Fig. 3, Figs. S5–7, S10**). During our reaction monitoring, we discovered a previously unreported activity of the Flexizyme, namely the ribozyme-catalyzed hydrolysis of aminoacyl-RNA esters in the presence of DBE-OH, the hydrolysis product of DBE-gly (**Fig. S5**). After accounting for this unusual ribozyme activity, as well as the reactions shown in **Fig. 3**, our *in silico* autocatalysis kinetic model faithfully reproduced the sigmoidal kinetics we observed experimentally (**Fig. 2B**, blue trace). Importantly, the trace is a kinetic prediction based on measuring individual rates and not a fit to the data.

**Fig. 3.**
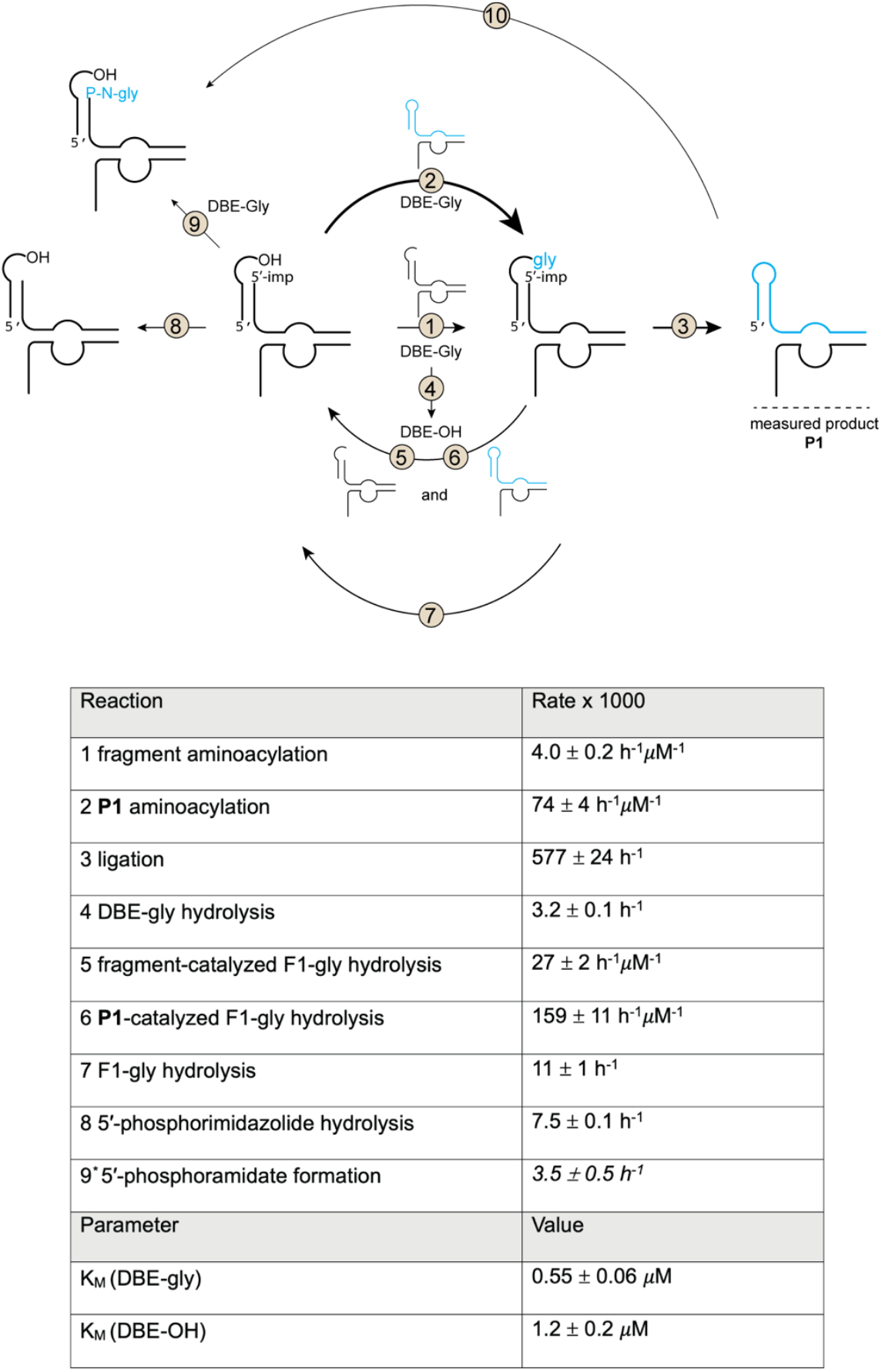
The autocatalytic reaction network. Top: the full network of reactions that were characterized in the autocatalytic P1 Flexizyme assembly network. The assembled, chimeric P1 ribozyme is represented in cyan. The arrow thickness corresponds roughly to the relative reaction rates. Bottom: rates of indicated reactions and Km values of indicated substrates. Reaction 9* rate was estimated with 20 µM Flexizyme fragments and was not used in the model. Reaction 10 rate was measured in ref. 43 and was insignificant for the autocatalytic network.

With a reasonable kinetic model established, we performed two key tests of whether the self-assembly process was autocatalytic or not. The first test involved performing the reaction in the absence of the catalytic enhancement by the product (i.e. the chimeric P1 Flexizyme). We performed this test with our kinetic model *in silico* because experimentally any changes that abolished the P1 activity also abolished the activity of the initial noncovalently assembled fragments. Computationally removing the added aminoacylation activity of the chimeric P1 Flexizyme resulted in an attenuated assembly yield and non-sigmoidal kinetics (**Fig. 2B**, orange trace). The second test for autocatalysis consisted of experimentally spiking in the reaction product at the beginning of the reaction and monitoring the increase in the self-assembly rate (**Fig. 4A**). Indeed, adding the covalently linked chimeric P1 product to our fragment system boosted the initial assembly kinetics and the overall reaction yield, an effect that could not be reproduced by simply increasing the concentration of fragments (**Fig. 4B**, circles). Crucially, the predictions of our kinetic model again matched the experimental data (**Fig. 4B**, traces).

**Fig. 4.**
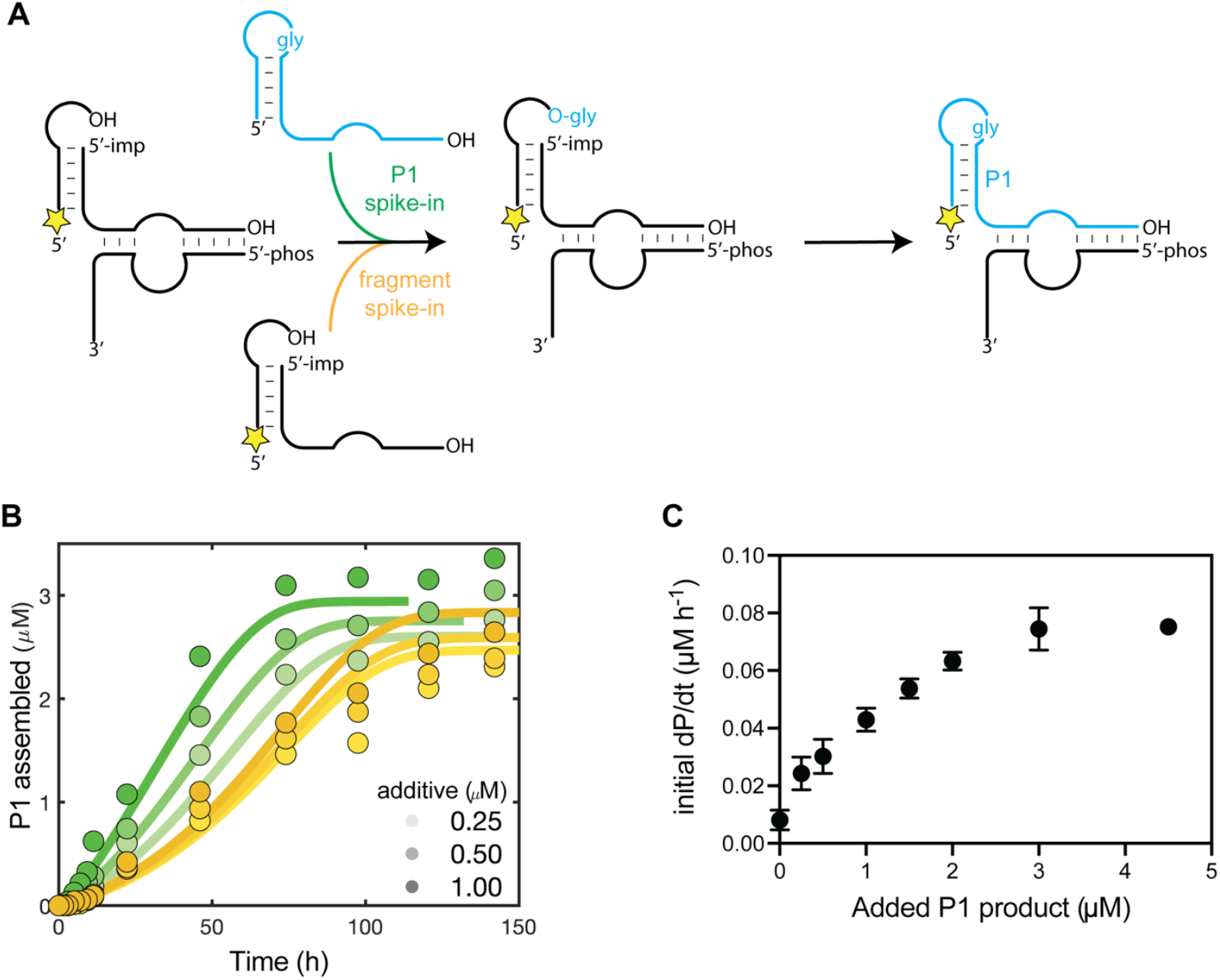
The effect of spiking in the chimeric P1 Flexizyme or the fragments on the autocatalytic cycle. **A:** General scheme of the autocatalytic cycle with either P1 (green arrow) or fragment (yellow arrow) spike-in. Yellow star: 5′-FAM label. 5′-imp represents the 5′-phosphorimidazolide. Note that the spiked in P1 product was not 5′-FAM labeled. **B:** Assembly of P1 over time with 5 µM of 5′-FAM labeled fragment 1, 5 µM of 5′-phosphorimidazolide fragment 2, and 0.6 µM of fragment 3, with the spike-in of either chimeric P1 (green) or fragments (yellow) at the additive concentrations of 0.25, 0.5, and 1 µM. The reaction was followed by acidic denaturing PAGE. The filled circles represent the collected data. The traces represent the predicted reaction timecourse from the global kinetic model. **C:** The initial reaction rates (dP/dt; P being the ligated P1 chimeric Flexizyme) plotted versus the added chimeric P1 product. See Fig. S8 for the initial rate data.

Adding increasing concentrations of the covalently linked chimeric P1 product to the reaction mix allowed us to determine the reaction order with respect to the catalyst, i.e. the P1 product, by plotting the initial reaction rate versus the concentration of the spiked-in catalyst. We found that the reaction rate increased nearly linearly with increasing intermediate catalyst concentrations (**Figs. 4C and S8**), implying a reaction order of close to 1 in catalyst, consistent with exponential growth(*47*). At higher concentrations of catalyst, the initial rate plateaus, likely due to saturation of the binding of the P1 product to fragment 3, which is the limiting reagent (0.6 µM) in the reaction, and thus limits the maximum amount of active Flexizyme that can be assembled. Taken together, these results are consistent with the proposed autocatalytic assembly cycle shown in Fig.s 2A and S3 and inconsistent with a simple multi-step assembly process.

An attractive feature of autocatalytic RNA reactions is their potential ability to sustain themselves indefinitely given the addition of fresh substrates, once the reaction is initiated. Templated ligation reactions, with the exception of the cross-replicating amd cross-chiral ribozyme ligases(*23, 48*), cannot sustain themselves indefinitely due to product inhibition. To test whether the autocatalytic P1 Flexizyme can sustain itself indefinitely, we allowed the autocatalytic reaction to self-initiate for 72 hours before diluting it four-fold into a freshly prepared uninitiated reaction mixture. We repeated the four-fold dilution every 72 hours for a total of 648 hours. Despite the dilution, the P1 Flexizyme assembly reaction accelerated until reaching a plateau after four cycles of dilution and assembly (**Fig. 5**, black trace). With each subsequent dilution, the concentration of the newly synthesized P1 Flexizyme increased until it matched the concentration present before dilution. In parallel, we also performed the same serial dilution experiment with a reaction that was seeded with 0.5 µM of the preformed P1 Flexizyme at the outset. This reaction reached steady state kinetics after only one dilution, but its steady state yield was the same as that of the unseeded reaction (**Fig. 5**, orange trace), suggesting that the advantage of seeding the assembly reaction with the preformed P1 Flexizyme becomes erased relatively quickly. The only other examples of self-sustained RNA assembly with cross-replicating or cross-chiral ligase ribozymes began by seeding the reaction with the preformed catalyst to initiate the reaction. In contrast, our results are consistent with a self-sustained autocatalytic RNA assembly process that is self-initiated.

**Fig. 5.**
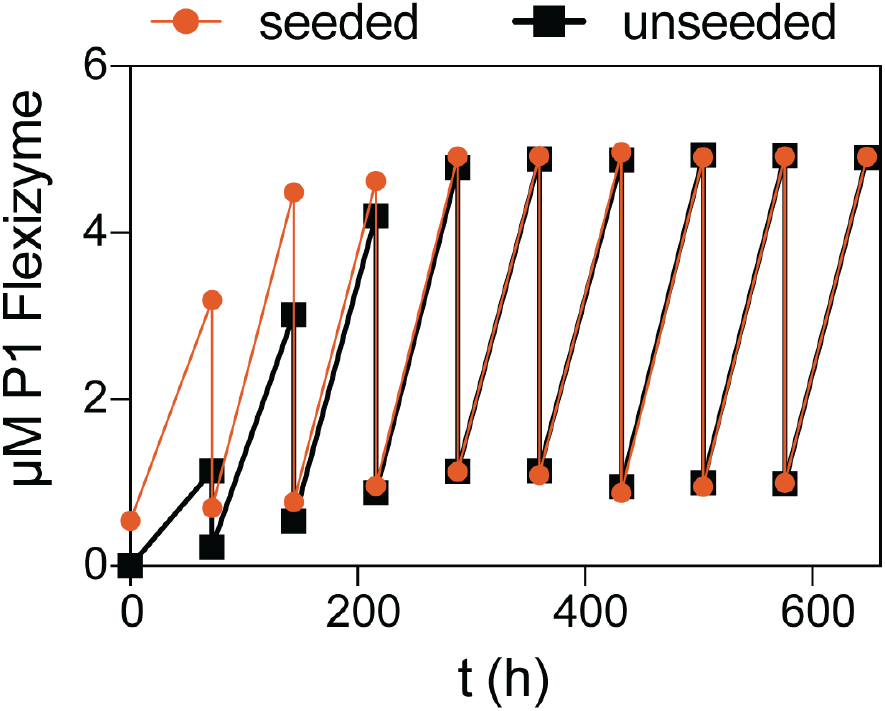
Self-sustained assembly of the chimeric P1 Flexizyme. Four-fold serial dilution of the autocatalytic P1 Flexizyme assembly reaction into a fresh, uninitiated assembly reaction was performed every 72 hours for 648 hours (black trace). The same procedure was repeated for a reaction that was seeded with 0.5 µM of the preformed P1 Flexizyme (orange trace). Reaction conditions: 0 ºC, 100 mM imidazole pH 8, 5 mM MgCl_2_, 5 µM of 5′-FAM labeled fragment 1, 5 µM of 5′-phosphorimidazolide fragment 2, 0.6 µM of fragment 3, and 3.38 mM DBE-gly.

Because the chimeric Flexizyme aminoacylates RNA substrates *in trans*, it could potentially also facilitate the assembly of other chimeric ribozymes with unrelated functions. To test this possibility, we incubated the chimeric P1 Flexizyme with two RNA fragments derived from the hammerhead ribozyme (**Fig. S9A**). We were gratified to observe rapid assembly of the chimeric hammerhead, in a reaction that was dependent on the presence of the chimeric Flexizyme (**Figs. 6A, S9B**). To test the activity of the newly assembled chimeric hammerhead, we diluted the assembly reaction without any purification into the hammerhead reaction buffer containing its substrate. The chimeric hammerhead displayed robust substrate cleavage activity that was substantially higher than the activity of the two noncovalently assembled hammerhead fragments but lower than the full-length, all-RNA hammerhead (**Figs. 6, S9B**). This result demonstrates that the autocatalytic P1 Flexizyme can not only assemble itself but can also assemble other catalytic chimeric RNAs.

**Fig. 6.**
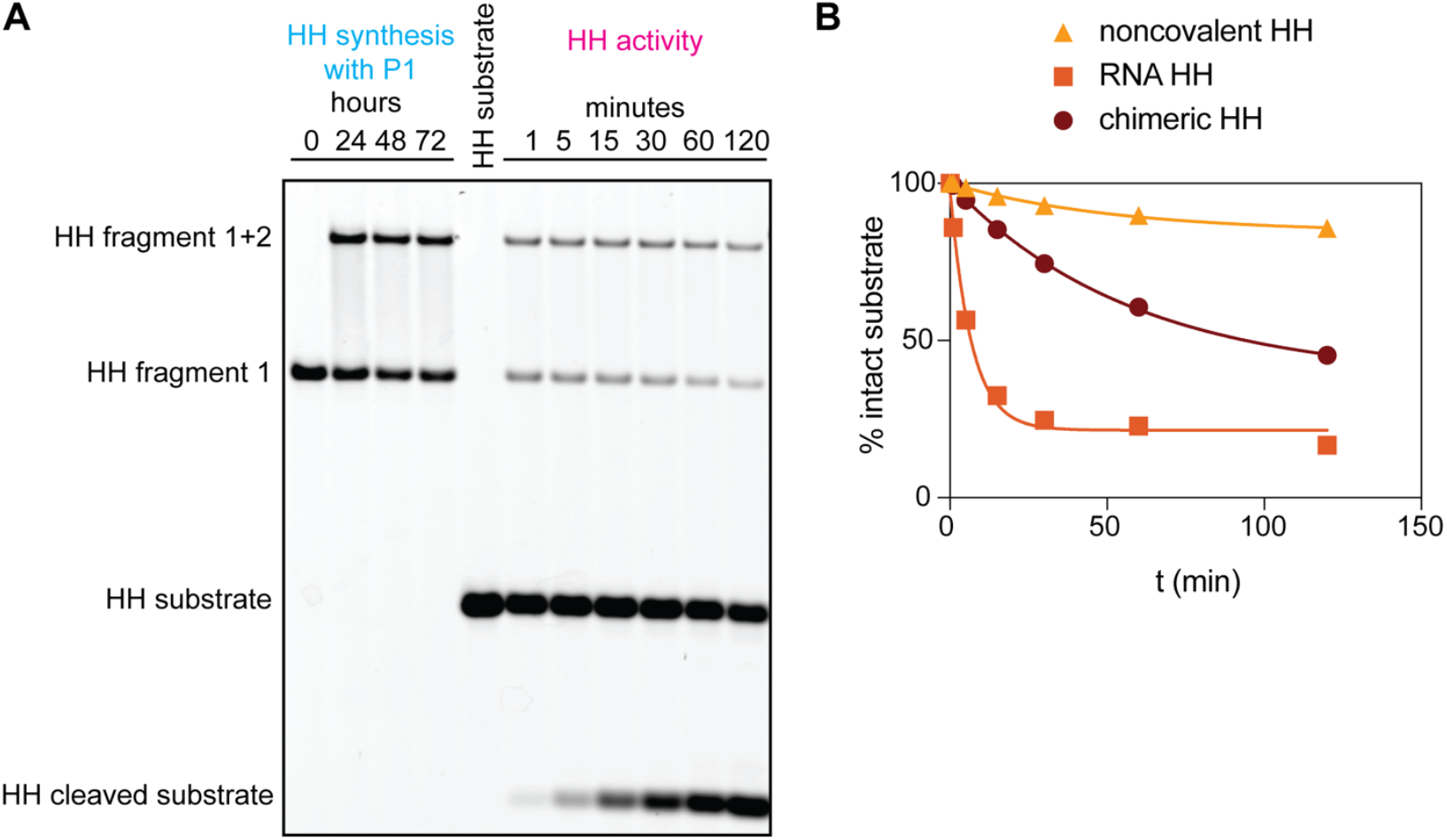
Assembly of an active, chimeric hammerhead by the chimeric P1 Flexizyme. **A:** Denaturing PAGE of the P1-catalyzed chimeric hammerhead assembly and activity. Conditions for HH synthesis: 5 mM MgCl_2_, 100 mM imidazole pH 8, 2 µM chimeric P1 Flexizyme, 3.38 mM DBE-gly, 5 µM 5′-FAM labeled HH fragment 1, and 5 µM of HH 5′-phosphorimidazolide fragment 2. Conditions for HH activity: 37 ºC, 5 mM MgCl_2_, 100 mM HEPES pH 7, 0.2 µM HH substrate. The assembly reaction was diluted 100-fold into the HH activity buffer to initiate the reaction. **B:** Time course of substrate cleavage by the noncovalently assembled hammerhead, the full-length all-RNA hammerhead, and the chimeric hammerhead. The lines are not fits but are intended to guide the eye. See also Fig. S9.

## Discussion

Our work demonstrates that the autocatalytic self-assembly of ribozymes can be driven by aminoacylation followed by loop-closing ligation. Most previously reported autocatalytic ribozymes rely on sequence-defined templated ligation to drive assembly, which eventually leads to the accumulation of non-productive base-paired duplexes, such that self-assembly becomes off-rate llimited. We circumvent this limitation by using aminoacylation coupled with loop-closing ligation, a process that buries the aminoacylated RNA substrate in a highly structured product stem-loop(*43*). The formation of the structured loop prevents ribozyme-substrate reassociation, thus avoiding product inhibition of the catalyst. This non-templated nature of loop-closing ligation allowed us to demonstrate uninhibited, self-sustained assembly of the chimeric glycine-bridged Flexizyme, which can proceed indefinitely over cycles of serial dilution into fresh substrates. Autocatalytic self-assembly of the *Azoarcus* group I intron from its fragments likely also avoids product inhibition by structurally burying the nucleotides involved in base-pairing with the catalyst after the ligation step is completed(*24*). A more recent example of uninhibited autocatalytic RNA assembly features two cross-chiral ligases that were evolved *in-vitro* to facilitate multi-turnover ligation reactions, thus avoiding the product-catalyst inhibition and sustaining each other indefinitely(*48*). Our autocatalytic chimeric Flexizyme has the added advantage that, in addition to being able to self-assemble, it can also assist in the assembly of other chimeric RNA stem-loops if they contain appropriate recognition overhangs. We demonstrate this advantage by using the self-assembled chimeric Flexizyme to assemble a chimeric hammerhead ribozyme through the same process of aminoacylation followed by loop-closing ligation.

We have developed a kinetic model of the chimeric Flexizyme assembly reaction network that accurately predicts the self-assembly process over a range of RNA concentrations. We expect that further optimization of this kinetic model will allow for exploration of the potential for autocatalytic self-assembly under more complex and realistic conditions, for example in the presence of complex mixtures of RNA oligonucleotides. Incorporating the binding and inhibition parameters of the additional oligonucleotides expected to be present in a self-replicating protocell could lead to more realistic reaction networks directly relevant to the origin of life(*26, 49, 50*). This could lead to the exciting prospect of testing whether autocatalytic RNA replication cycles, such as the virtual circular genome(*51*), could not only generate the necessary longer RNA oligonucleotides but also enable their assembly into active ribozymes. We suggest that the autocatalytic assembly of an aminoacyl-RNA synthetase could facilitate the emergence of diverse ribozymes including ribozymes that enhance RNA replication. Generation of multiple useful ribozymes could help to explain how a self-sustaining genotype could encode multiple self-sustaining phenotypes.

We have described a potentially primordial scenario that would have conferred a fitness advantage to RNAs that participated in aminoacylation, either as substrates or catalysts. In our simple reaction scheme, three relatively short oligonucleotides catalyze aminoacylation of one of the component oligonucleotides, which after loop-closing assembly generates a much more efficient RNA aminoacylation catalyst. Indefinite propagation of this catalyst, given a supply of fresh substrates, ensures the preservation of the RNA aminoacylation phenotype by coupling it to self-replication of the RNA aminoacylation catalyst. Importantly, the aminoacyl-RNA synthetase ribozyme is capable of catalyzing the assembly of other chimeric ribozymes that catalyze unrelated reactions. A general catalyst of RNA aminoacylation (such as the Flexizyme) could incorporate amino acids with diverse side chains into chimeric ribozymes, potentially leading to the emergence of catalysts whose activity depended on the identity of the incorporated amino acid. Such a scenario would create a selection pressure to place the correct amino acid in the correct RNA sequence context, favoring the evolution of specialized aminoacyl-RNA synthetase ribozymes with RNA- and amino acid-specificity. This sequence of events could potentially explain how the RNA World generated specific and diverse aminoacyl-RNA substrates and synthetases, the essential ingredients for the evolution of translation, without invoking translation itself as the selection pressure.

## Acknowledgments

The authors are grateful to Profs Donna Blackmond and Gerald Joyce for thoughtful discussions and the Szostak laboratory for their suggestions on the work.

## Funding

JWS is an investigator of the Howard Hughes Medical Institute (JWS) Simons Foundation Grant 290363 (JWS)

## Author contributions

Conceptualization: AR, JWS

Methodology: AR, MT, AM, JWS

Investigation: AR

Visualization: AR, MT

Funding acquisition: JWS

Project administration: AR, JWS

Supervision: AR, JWS

Writing: AR, JWS

## Competing interests

Authors declare that they have no competing interests.

## Data and materials availability

All data and code are available upon request.

## Supplementary Materials

Materials and Methods Figs. S1 to S10

Table S1

References (*52, 53*)

## Notes

### Competing Interest Statement

The authors have declared no competing interest.

