## Supplementary Information for "Autocatalytic assembly of a chimeric aminoacyl-RNA synthetase ribozyme"

**The PDF file includes:**

Materials and Methods
Figs. S1 to S10
Table S1

### Materials and Methods

General information.

All reagents were purchased from Sigma-Aldrich and Fisher Scientific, unless otherwise noted. Oligonucleotide synthesis reagents were purchased from ChemGenes and Glen Research.

Oligonucleotide synthesis.

All oligonucleotides were synthesized in-house on a K&A H-6 instrument. Coupling times were adjusted according to the manufacturer's instructions (Glen Research or ChemGenes). Oligonucleotides were cleaved and the nucleobases deprotected as recommended by the manufacturer. The 2′-OTBDMS protecting groups were removed by dissolving the oligonucleotides in 100 µL dimethylsulfoxide and 125 µL triethylamine trihydrofluoride and incubating them at 65 ºC for 2.5 hours. Following precipitation with 0.1 volumes of 5 M ammonium acetate and 1 mL isopropanol and one wash with 80 % aq ethanol (v/v), the oligonucleotides were dissolved in 99 % formamide (v/v) 5 mM EDTA and purified by denaturing PAGE. The desired gel bands were isolated, crushed, and soaked in a solution of 5 mM sodium acetate pH 5.5 and 2 mM EDTA for 16 hours. The extracted oligonucleotides were desalted using C18 Sep-Pak cartridges (Waters).

Oligonucleotide activation.

To a 200 µL solution of 5′-phosphorylated oligonucleotides in imidazole pH 7 buffer was added EDC.HCl so that the final concentrations were 200 µM oligonucleotide, 100 mM imidazole pH 7, and 100 mM EDC.HCl. The solution was incubated at room temperature for 2 hours before precipitation with 0.1 volumes of sodium perchlorate-saturated acetone and 1 mL of cold acetone. After two washes with cold 1:1 acetone:diethylether (v/v), the pellet was dried and redissolved in 1 mM imidazole pH 8 buffer. The activated oligonucleotides were stored at -80 ºC until use.

Aminoacylation reactions.

1. Reaction with Flexizyme **fragments**: To a solution containing the Flexizyme fragments in imidazole pH 8 buffer was added $MgCl_2$ and the dinitrobenzyl ester of glycine (DBE-gly) so that the final concentrations were: 5 mM $MgCl_2$, 100 mM imidazole pH 8, DBE-gly (see Fig. captions for concentrations), fragments (see Fig. captions for concentrations). Note that fragment 1 is the fluorescently labeled Flexizyme substrate that is used to measure the aminoacylation activity.
2. Reaction with the covalently linked Flexizyme **P1**: To a solution containing the covalently linked Flexizyme P1, fragment 3, the fluorescently labeled fragment 1, and the partial complement of the fragment 1 in imidazole pH 8 buffer was added $MgCl_2$ and the dinitrobenzyl ester of glycine (DBE-gly) so that the final concentrations were: 5 mM $MgCl_2$, 100 mM imidazole pH 8, DBE-gly (see Fig. captions for concentrations), 0.5 µM P1 Flexizyme, 5 µM fragment 1 substrate, and fragment 3 (see Fig. captions for concentrations). Note that fragment 1 is the fluorescently labeled Flexizyme substrate that is used to measure the aminoacylation activity.
3. Reaction with the covalently linked Flexizyme **P1+P2**: To a solution containing the covalently linked Flexizyme P1+P2 and the fluorescently labeled fragment 1 in imidazole pH 8 buffer was added $MgCl_2$ and the dinitrobenzyl ester of glycine (DBE-gly) so that the final concentrations were: 5 mM $MgCl_2$, 100 mM imidazole pH 8, DBE-gly (see Fig. captions for concentrations), 0.5 µM P1+P2 Flexizyme, and 5 µM fragment 1 substrate.

Aminoacylation reactions were incubated at 0 ºC, with 1 µL aliquots quenched in 4 µL of the acidic quenching buffer (10 mM EDTA pH 8.0, 1x bromophenol blue, 100 mM sodium acetate pH 5.0, 150 mM HCl, 75 % v/v formamide) prior to loading 2.5 µL of the quenched reactions into acidic 20 % denaturing polyacrylamide gels. The acidic gels were cast using 100 mM sodium acetate pH 5 instead of 1x Tris-Borate-EDTA buffer. The acidic PAGE was performed at 4 ºC at 25 W for 3 hours. The gels were scanned in an Amersham Typhoon (Cytiva) gel imager and the bands were analyzed using ImageQuant TL software. Aminoacylation percentages were obtained by dividing the intensity of the aminoacylated band by the sum of the intensities of the aminoacylated and nonaminoacylated bands for each lane.

<u>Chimeric Flexizyme synthesis for aminoacylation and spike-in studies.</u>
**P1+P2** Flexizyme: Fragments corresponding to this ribozyme (see Fig. S1) with fragments 2 and 3 activated as 5′-phosphorimidazolides, were added to a solution of imidazole pH 8, $MgCl_2$, and DBE-gly. Concentrations in the 500 µL total volume were: 5 µM of each fragment, 100 mM imidazole pH 8, 5 mM $MgCl_2$, and 3.38 mM DBE-gly. The reaction was incubated at 0 ºC for 19 hours, before being concentrated to 50 µL using 0.5 mL 10k MWCO Amicon Filters. The concentrated reactions were mixed with 50 µL of neat formamide and purified on 16 % denaturing PAGE. The desired bands were cut out, crushed, and soaked for 3 hours at 4 ºC in 5 mM sodium acetate, 2 mM EDTA acidified to pH 5 with 1 M HCl. The extracted RNA was concentrated with 0.5 mL 10k MWCO Amicon Filters and desalted using the Zymo RNA Clean & Concentrator kit, according to the manufacturer protocol.
**P1** Flexizyme (see Fig. S1): The assembly reaction was set up as above, except that fragment 3 was not activated as 5′-phosphorimidazolide and its final concentration was 1 µM instead of 5 µM. The reaction was incubated at 0 ºC for 96 hours before being purified and desalted as above.

<u>Aminoacyl-RNA hydrolysis reactions.</u>
1. Preparative synthesis of the glycylated fragment 1: A 500 µL reaction containing 100 mM imidazole pH 8, 5 mM $MgCl_2$, 10 µM fragment 1, 1 µM dFx_S7 Flexizyme (Table S1), and 3.38 mM DBE-gly was allowed to proceed for 48 hours at 0 ºC. The reaction was concentrated using 0.5 mL 3k MWCO Amicon Filters to 50 µL, diluted with 50 µL of formamide, and purified by preparative acidic denaturing PAGE. The aminoacylated fragment 1 band was cut out, crushed, and soaked in 5 mM sodium acetate, 2 mM EDTA adjusted to pH 5 for 3 hours at 4 ºC. The supernatant was filtered, concentrated with 3k MWCO Amicon Filters to 50 µL, and precipitated with 5 µL of 3 M sodium acetate pH 5 and 1 mL cold ethanol. Following two washes with 80 % v/v ethanol, the material was analyzed by analytical acidic PAGE to obtain the aminoacylated fraction.
2. The material prepared above was then subjected to hydrolysis under the same conditions as the aminoacylation reaction without the addition of DBE-gly.
3. The deacylation activity of the Flexizyme was characterized by adding DBE-OH, the hydrolysis product of DBE-gly, to the hydrolysis reaction in varying concentrations.

<u>Aminoacyl-RNA ligation reaction.</u>
A 10 µL reaction containing 100 mM imidazole pH 8, 5 mM $MgCl_2$, 5 µM glycylated fragment 1, 5 µM fragment 2 activated as a 5′-phosphorimidazolide, and 3.38 mM DBE-gly was incubated at 0 ºC. Aliquots (1 µL) were taken at various time points and quenched with 4 µL quenching buffer

(90 % v/v formamide, 20 mM EDTA, 1x bromophenol blue). The quenched aliquots were analyzed by 20 % denaturing PAGE and the % ligated product was obtained by dividing the intensity of the ligated band by the sum of the intensities of the ligated and unligated bands for each lane.

DBE-gly hydrolysis reaction.

A 500 µL reaction containing 100 mM imidazole pH 8 (prepared in $D_2O$), 5 mM $MgCl_2$, and 3.38 mM DBE-gly (prepared in d6-DMSO) was incubated at 0 ºC. $^1H$ NMR spectra were taken at the indicated time points. The % DBE-gly was obtained by dividing the integrated singlet signal of the two alpha protons of DBE-gly by the sum of the integrated alpha proton signal of DBE-gly and DBE-OH.

5′-phosphorimidazolide hydrolysis reaction.

A short model oligonucleotide based on fragment 2 (see Table S1) was designed in order to achieve baseline separation between the 5′-phosphorimidazolide activated oligonucleotide and the hydrolyzed 5′-phosphate oligonucleotide by analytical HPLC. The full-length fragment 2 could not be fully resolved to obtain accurate data for the kinetic model. We estimated the % initial activation of fragment 2, that we used to constrain the kinetic model of the overall assembly reaction, based on the maximum yield of the aminoacyl-RNA ligation reaction.

The 5′-phosphorimidazolide hydrolysis reaction contained 50 µL of 100 mM imidazole pH 8, 5 mM $MgCl_2$, 3.38 mM DBE-gly, 20 µM of fragment 1 treated with sodium periodate to prevent 2′,3′-diol aminoacylation and subsequent ligation, 20 µM of model oligonucleotide activated as a 5′-phosphorimidazolide, and 20 µM of dFx_S7 (see Table S1). Fragment 1 and dFx_S7 were included to control for any acceleration of the hydrolysis reaction by the Flexizyme. During the incubation at 0 ºC, 5 µL aliquots were taken and analyzed by HPLC using the Atlantis TM T3 column (3 µm, 4.6 x 150 mm) at a flow rate of 0.5 mL/min. The following gradient was used: (A) aqueous 50 mM triethylammonium acetate (pH 7.0) and (B) acetonitrile, from 6% to 12% B over 20 min. The % 5′-phosphorimidazolide remaining was determined by diving the area under the signal for the activated species by the sum of the signals for the activated and hydrolyzed oligonucleotide.

Sodium periodate oxidation of fragment 1.

A 500 µL reaction containing 5 µM fragment 1 and 10 mM sodium periodate was allowed to proceed for 1 hour at 0 ºC. The reaction was then concentrated using 3k MWCO Amicon Filters to 50 µL, followed by precipitation with 5 µL sodium acetate pH 5.5 and 1 mL cold ethanol. After two 80 % v/v ethanol washes, the pellet was dissolved in ultra-pure water and used without further purification.

Glycine-5′-phosphoramidate formation reaction.

A 100 µL reaction containing 100 mM imidazole pH 8, 5 mM $MgCl_2$, 3.38 mM DBE-gly, and 20 µM of 5′-phosphorimidazolide activated fragment 2 was incubated at 0 ºC. After 72 hours, a 5 µL aliquot was analyzed by analytical HPLC as above, resulting in no new chromatographically resolved signals. Another 5 µL aliquot at the 72 hours timepoint was analyzed on the Agilent 6540 mass spectrometer. Exact masses corresponding to three species were identified and shown in Fig. S7.

The overall assembly reaction of the chimeric Flexizyme from fragments.

The assembly reaction was set up at 0 ºC in a volume of 10 μL containing 5 μM of Flexizyme fragments 1 and 2, with the fragment 2 activated as 5′-phosphorimidazolide, 0.6 μM of fragment 3, 100 mM imidazole pH 8, 5 mM $MgCl_2$, and 3.38 mM DBE-gly. The reaction was incubated at 0 ºC, and 0.7 μL aliquots were quenched at various time points in acidic quenching buffer (10 mM EDTA pH 8.0, 1x bromophenol blue, 100 mM sodium acetate pH 5.0, 150 mM HCl, 75 % v/v formamide) prior to loading 2.5 μL of the quenched reactions into acidic 20 % denaturing polyacrylamide gels. The gels were run and analyzed as above.

The overall assembly reaction of the chimeric Flexizyme from fragments with the addition of preformed P1 Flexizyme or fragments was set up as above, except preformed, unlabeled P1 Flexizyme or the equivalent molar amount of fragments was added at 0.25, 0.5, and 1 μM.

Kinetic modeling of the overall assembly reaction.

The complex reaction network studied in this work was modeled with a series of coupled first-order differential equations accounting for the transformation of 8 species over time as described by the 8 chemical reactions in Fig. 3. Time traces were simulated by integrating the model using Euler's method. Binding of DBE-Gly and DBE-OH to the fragmented Flexizyme and **P1** were captured by simple hyperbolic curves for competitive binding updated at every integration time-step(*52*). Under the assumption of fast reshuffling of all short oligonucleotides constituting the Flexizymes over the reaction timescale, the concentrations of all possible complexes were updated at every integration time-step using a MATLAB implementation of the Thomas Wayne Wall algorithm for the solution of multiple chemical equilibria(*50, 53*). To parameterize and build the model, a set of experiments specifically designed to be maximally sensitive to different and complementary subsets of parameters were fitted either with simple exponentials or with simulated traces from the aforementioned model using MATLAB non-linear fit algorithm (*nlinfit*). Errors on the coefficients reported in this work have been derived from the provided variance-covariance matrix. A full list of fitted parameters and associated standard errors is presented in Fig. 3.

For simple hydrolysis reactions of labile species, such as reactions number 4 (DBE-gly hydrolysis to DBE-OH), number 7 (hydrolysis-mediated deacylation of fragment 1) and number 8 (5′-phosphorimidazolide hydrolysis to yield a non-reactive 5′-phosphorylated fragment 2), these datasets were individually fitted with a single exponential model with free amplitude to capture any offset in the initial purity of the compounds (see Supplementary Figs. 6B,C,D).

Given the complexity of the reaction network, no analytical solution for the concentration of species as a function of time is readily available and we opted to model it using a series of coupled first-order differential equations, integrated using Euler's method to generate time traces. The rates and parameters that could not be estimated independently had to be globally fitted using MATLAB *nlinfit* using simulated traces on data deriving from multiple experiments specifically designed to be maximally sensitive to different and complementary subsets of parameters (see Figs. S5 and S6A). Errors on the coefficients here reported have been directly calculated from the variance-covariance matrix provided by MATLAB.

We started to assemble the model based on our understanding as derived from established literature, so that the formation of P1 would be the outcome of the sequential reactions of aminoacylation of F1 and loop-closing of the F1-gly and 5′-phosphorimidazolide F2 complex.

Surprisingly, we found that aminoacylation experiments show a strong non-monotonic behavior over time, that we could not explain in terms of hydrolysis of the acylated fragment 1 substrate as independently determined for the hydrolysis-mediated deacylation of F1-gly ($k_7$). A systematic study of this behavior revealed that the drop in acylated products is dependent on

Flexizyme concentration, and is also dependent on DBE-OH concentration. We therefore hypothesized that this class of ribozymes effectively catalyzes the deacylation of F1-gly using DBE-OH, which accumulates over time due to DBE-gly hydrolysis. Indeed, we confirmed that the Flexizyme has this secondary catalytic activity, which has not been previously reported.

Measured apparent reaction rates for catalyzed aminoacylation ($k_1^*$ & $k_2^*$) and catalyzed deacylation ($k_5^*$ & $k_6^*$) were found to be dependent on DBE-gly and DBE-OH concentrations respectively. To capture this behavior in our model, the rates for these chemical reactions were updated at every integration step adapting to the relative fraction of Flexizyme bound to either DBE-gly or DBE-OH, so that the effective Flexizyme-catalyzed acylation rates would be:

$$k_1^* = k_1 * f_{DBE-gly}$$
$$k_2^* = k_2 * f_{DBE-gly}$$

And the effective Flexizyme-catalyzed deacylation rates would be:

$$k_5^* = k_5 * f_{DBE-OH}$$
$$k_6^* = k_6 * f_{DBE-OH}$$

The fraction of Flexizyme bound to DBE-gly ($f_{DBE-gly}$) and the fraction of ribozyme bound to DBE-OH ($f_{DBE-OH}$) were approximated following a simple competitive binding model under the assumption of a common binding site:

$$f_{DBE-gly} = \frac{[DBE-gly]}{K_{DBE-gly}\left(1 + \frac{[DBE-OH]}{K_{DBE-OH}}\right) + [DBE-gly]}$$
$$f_{DBE-OH} = \frac{[DBE-OH]}{K_{DBE-OH}\left(1 + \frac{[DBE-gly]}{K_{DBE-gly}}\right) + [DBE-OH]}$$

In this sense, $k_1$, $k_2$, $k_5$ and $k_6$ reported in this work can be effectively considered as maximum rates in the limit of Flexizyme-saturated either by DBE-gly ($k_1$, $k_2$) or DBE-OH ($k_5$, $k_6$). Interestingly, the binding constants for DBE-gly and DBE-OH were found to be comparable, supporting the previously established hypothesis that the aromatic moiety is interacting with the Flexizyme binding site, and not the amino acid(*46*).

Finally, we had to account for the fact that the system is composed of species taking part in multiple equilibria. For example, F1 can be bound to either F2 or F2-im.
Given the large combinatorics raising from having multiple hybridizing RNA strands, we had to employ an automated method to update the distribution of species into different complexes. Under the assumption of (i) fast reshuffling of the short oligonucleotides constituting the fragmented Flexizyme over the reaction timescale(*52*) and (ii) strong binding (K << [F1],[F2],[F1-gly],…), the concentrations of all possible complexes could be updated at every integration time-step by solving a large system of equations describing all chemical equilibria and mass conservations. Unfortunately, the built-in *vpasolve* function in MATLAB was too slow to be practically used to solve such a large system and to yield traces usable for globally fitting our large dataset, so we opted for using a custom MATLAB implementation of the very efficient algorithm by Thomas Wayne Wall for the solution of multiple chemical equilibria(*50, 53*), with a reduction of execution times by two orders of magnitude.

Accounting for all the reactions here described, the overall autocatalytic reaction can be finally modeled as follows:

$$\frac{d[F1]}{dt} = -[F1][R]k_1^* - [F1][P1{:}F3]k_2^* + [F1-gly][R]k_5^* + [F1-Gly][P1{:}F3]k_6^* + [F1-gly]k_7$$

$$\frac{d[F1-gly]}{dt} = [F1][R]k_1^* + [F1][P1{:}F3]k_2^* - [F1-gly][R]k_5^* - [F1-gly][P1{:}F3]k_6^* - [F1-gly]k_7 - [F1-Gly{:}F2-im]k_3$$

$$\frac{d[F2]}{dt} = [F2-im]k_8$$

$$\frac{d[F2-im]}{dt} = -[F2-im]k_8 - [F1-gly{:}F2-im]k_3$$

$$\frac{d[P1]}{dt} = [F1-gly{:}F2-im]k_3$$

$$\frac{d[DBE-gly]}{dt} = -[DBE-gly]k_4$$

$$\frac{d[DBE-OH]}{dt} = [DBE-gly]k_4$$

Where R includes all non-covalent Flexizymes formed by a combination of F1, F1-gly, F2, F2-im and F3, namely F1:F2:F3, F1-gly:F2:F3, F1:F2-im:F3 and F1-gly:F2-im:F3.

It is important to note that the continuous lines reported in Fig. 2 of the main text are predictions and not fits. We found that the model at later times overestimates the reaction yield. We were able to identify a side reaction leading to the formation of a phosphoramidate species from 5′-phosphorimidazolide of fragment 2 and
DBE-gly through Mass Spectrometry (Fig. S7). Unfortunately, given that the reaction is dependent both on Flexizyme concentration and DBE-gly, we could not measure it outside of the autocatalytic cycle without using proxy oligonucleotides. To estimate the rate associated with this process, we performed a measurement using 20 μM of Flexyzime fragments 1 and 2, where the F1 fragment was periodate-treated (preventing it from being acylated) (Fig. S10), leading to an estimate of $k_9$ and suggesting that at least part of the overestimation could indeed be due to this side reaction.

Self-sustained assembly of the chimeric P1 Flexizyme.

Unseeded reaction: The assembly reaction was set up at 0 ºC in a volume of 10 μL containing 5 μM of Flexizyme fragments 1 and 2, with fragment 2 activated as a 5′-phosphorimidazolide, 0.6 μM of fragment 3, 100 mM imidazole pH 8, 5 mM $MgCl_2$, and 3.38 mM DBE-gly. The reaction was incubated at 0 ºC for 72 hours after which 2.5 μL of the reaction were mixed with 1 μL of a solution of 6 μM fragment 3 before being added to 6.5 μL of the fresh uninitiated reaction, so that the final volume was 10 μL and final concentrations of the reaction were one quarter of all of the reagents from the previous reaction, 5 μM of Flexizyme fragments 1 and 2, with the fragment 2 activated as 5′-phosphorimidazolide, 0.6 μM of fragment 3, 100 mM imidazole pH 8, 5 mM $MgCl_2$, and 3.38 mM DBE-gly. The four-fold dilution process was then repeated in the same manner seven more times.

Seeded reaction: The reaction was performed as above except the initial reaction contained 5 μM of Flexizyme fragments 1 and 2 and 0.5 μM of the preformed chimeric P1 product.

To monitor the reaction, 1 µL aliquots were quenched at the initiation, dilution, and completion of the reaction in acidic quenching buffer (10 mM EDTA pH 8.0, 1x bromophenol blue, 100 mM sodium acetate pH 5.0, 150 mM HCl, 75 % v/v formamide) prior to loading 2.5 µL of the quenched reactions into 20 % denaturing polyacrylamide gels. The gels were run and analyzed as above.

Chimeric hammerhead assembly and activity.

The assembly reaction was set up at 0 ºC in a volume of 10 µL containing 2 µM of unlabeled chimeric Flexizyme P1, 2 µM of Flexizyme fragment 3, 5 µM each of HH fragments 1 and 2, 100 mM imidazole pH 8, 5 mM $MgCl_2$, and 3.38 mM DBE-gly. The reaction was incubated at 0 ºC, and 0.7 µL aliquots were quenched at 0, 24, 48, and 72 hour time points in 4.3 µL of quenching buffer (90 % v/v formamide, 20 mM EDTA, 1x bromophenol blue).

At the 72 hour time point, 1 µL of the assembly reaction was diluted 5x with 4 µL of water. After dilution, 1 µL of the diluted assembly reaction was added to 19 µL of the hammerhead activity reaction with the final concentrations being 100 mM HEPES pH 7, 5 mM $MgCl_2$, and 0.2 µM HH substrate. For the RNA HH control reaction, the full-length RNA hammerhead was at 0.05 µM. The hammerhead reaction was then incubated at 37 ºC and 2 µL aliquots were quenched at 1, 5, 15, 30, 60, and 120 minute time points in 8 µL of quenching buffer (90 % v/v formamide, 20 mM EDTA, 1x bromophenol blue). The quenched aliquots were analyzed by 20 % denaturing PAGE as described above.

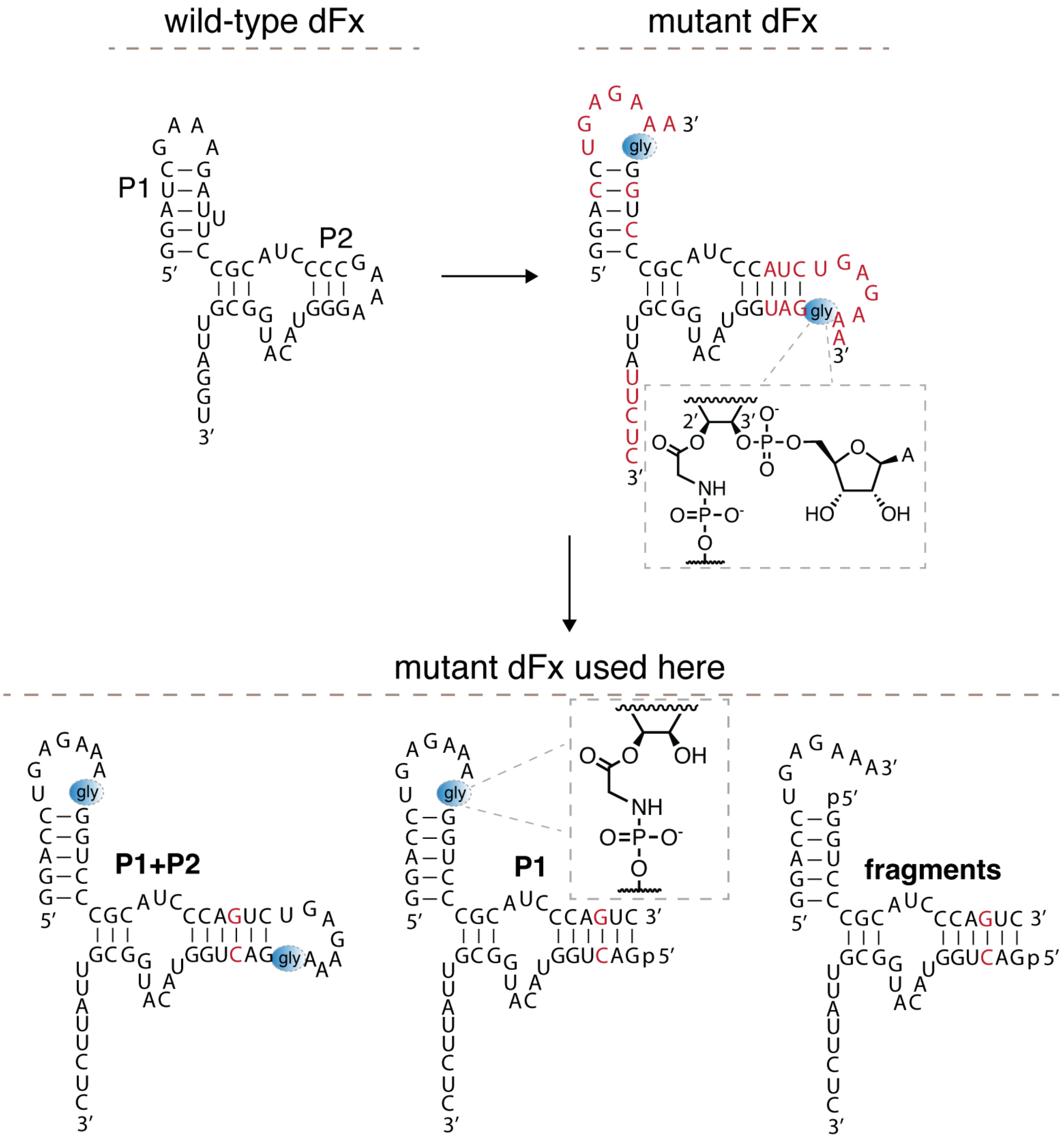


**Fig. S1. Alterations of the dFx Flexizyme.** Left: The secondary structure of the wild-type dFx Flexizyme. Right: The secondary structure of the dFx with the altered P1 stem, the extended P2 stem, the added 5′-UGAGAAA overhangs on the P1 and P2 stems, and the altered 3′-substrate binding site (modifications in red). The blue circles with "gly" represent the glycine bridges as prepared in ref. 43. Bottom: The secondary structures of the three Flexizyme constructs used in this study. The new modifications are indicated in red. Note that the glycine bridges are between the terminal 3′-nucleotides of the overhangs and 5′-phosphates, due to their Flexizyme-mediated preparation, which exclusively aminoacylates the 2′,3′-diol.

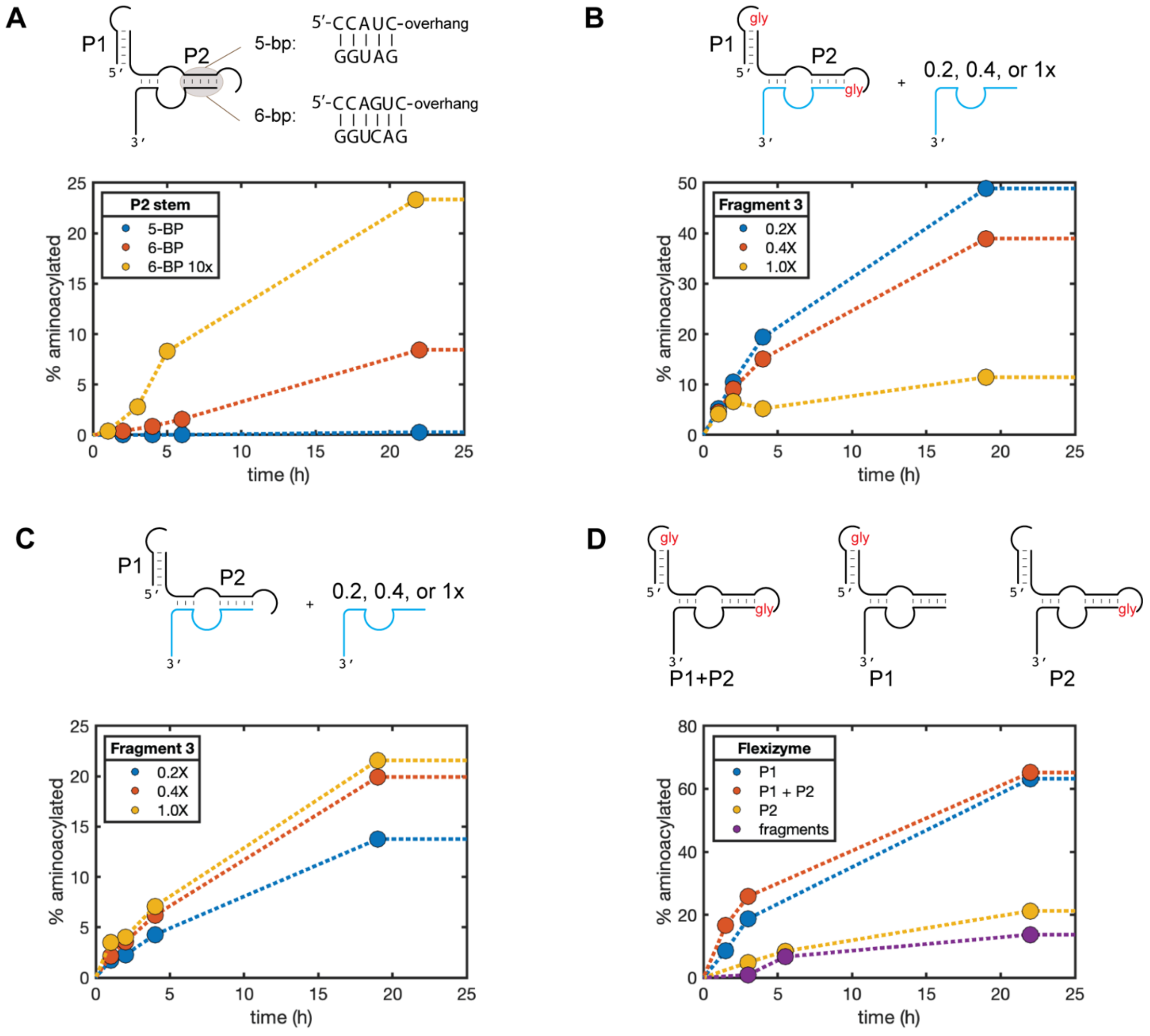


**Fig. S2. Aminoacylation optimization with the various fragmented and covalently linked Flexizyme constructs. A:** Aminoacylation activity of the Flexizyme fragments where the P2 stem contains 5 or 6 base-pairs was measured. The reactions contained 5 μM of the 5′-FAM fragment 1, 0.5 μM of fragment 2, and 0.5 μM of fragment 3, and 3.38 mM of DBE-gly. The 6-BP 10x condition contained 5 μM each of fragments 2 and 3. **B:** Aminoacylation activity of the fully covalently linked **P1+P2** Flexizyme in the presence of increasing concentration of fragment 3 was measured. The reactions contained 5 μM 5′-FAM labeled fragment 1, 5 μM fragment 2, 1 μM **P1+P2**, 3.38 mM DBE-gly, and either 1 μM, 2 μM or 5 μM of fragment 3 (denoted by 0.2X, 0.4X, and 1X, respectively). The third fragment significantly inhibited the activity of the covalent ribozyme, likely due to occupying the same binding site on the substrate fragment. Dilution of the third fragment relieves this inhibition. **C:** As in **B**, except no **P1+P2** was added. Dilution of the third fragment has minimal impact on the aminoacylation activity of the fragments. **D:** Aminoacylation activity was measured as in **B**, except fragment 3 was kept at the constant 1 μM. The red "gly" labels represent the glycine bridges. The traces in all four panels connect the data points for ease of visualization and do not correspond to fitted reaction rates.

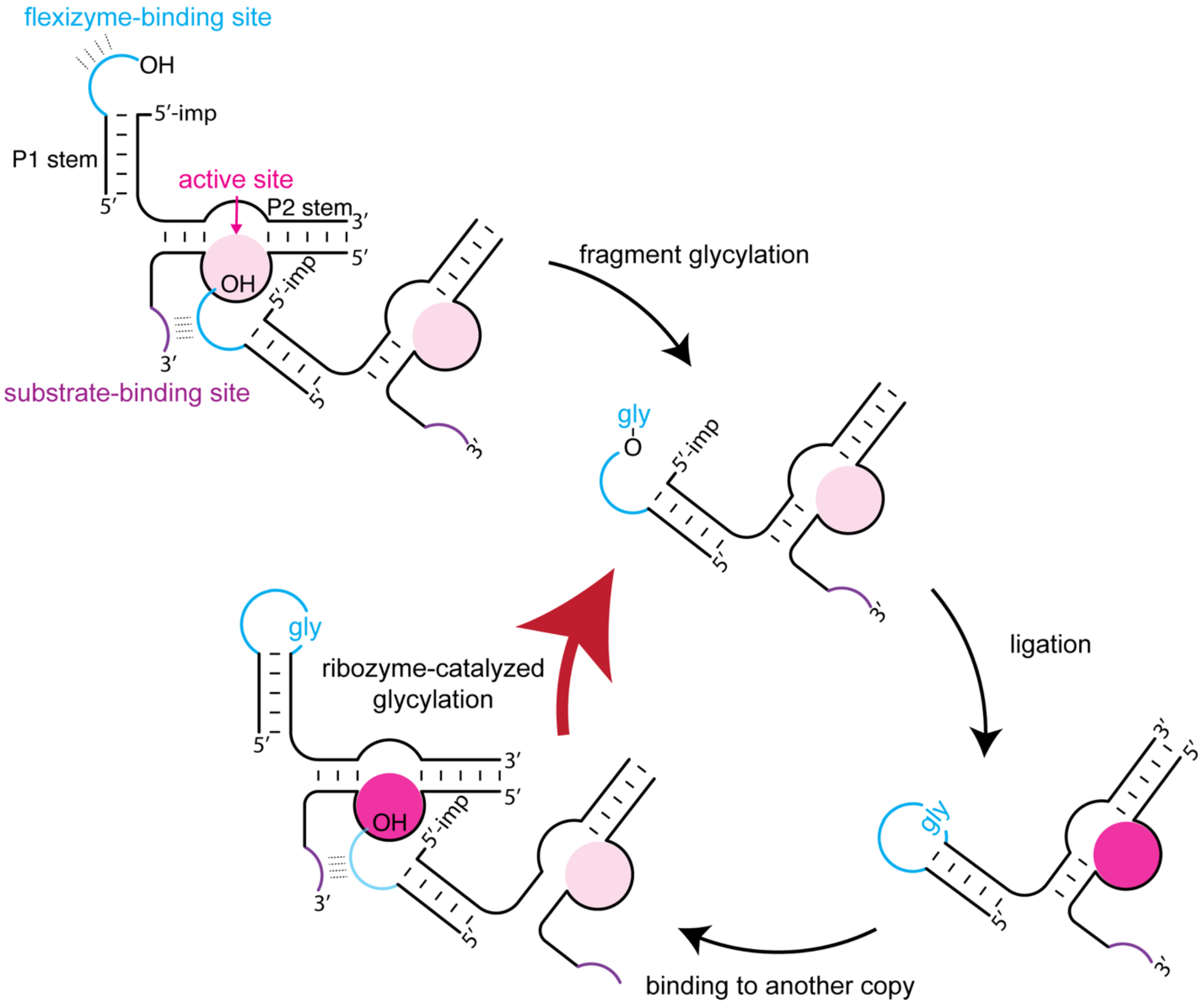


**Fig. S3. Autocatalytic assembly of the chimeric P1 Flexizyme shown in detail.** The chimeric Flexizyme ribozyme assembly according to the simplified scheme shown in Fig. 2. The overhang in light blue is UGAGAAA-3′, a sequence that can be bound by the purple UUCUC-3′ overhang of the Flexizyme. The active site for aminoacylation is depicted as a pink circle. The low level of activity of the fragments (active site in light pink) is sufficient to aminoacylate another copy of the fragmented ribozyme, which upon loop-closing ligation becomes more active (active site in dark pink) and facilitates its own assembly.

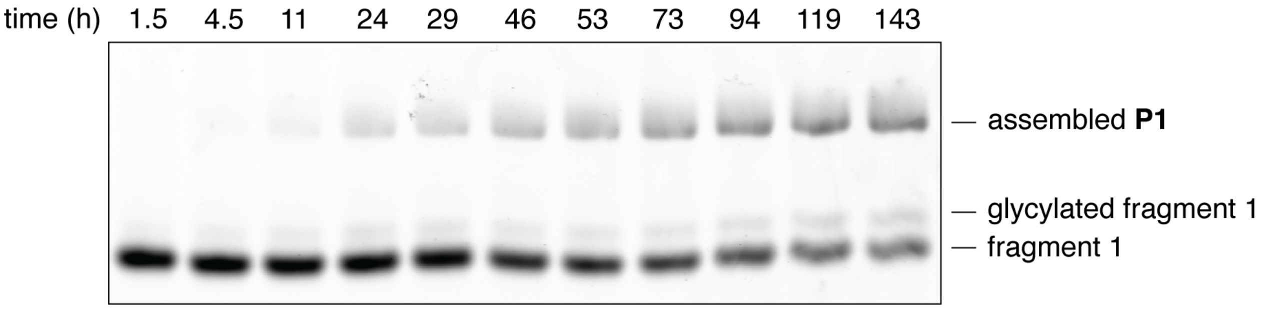


**Fig. S4. Representative gel image of the P1 Flexizyme assembly reaction.** The **P1** Flexizyme assembly reaction was prepared as described in the Methods and Fig. 2A and followed by acidic gel electrophoresis. The bottom-most band represents the 5′-FAM labeled fragment 1. The band just above it represents the glycylated fragment 1. The top-most band represents the assembled **P1** Flexizyme.

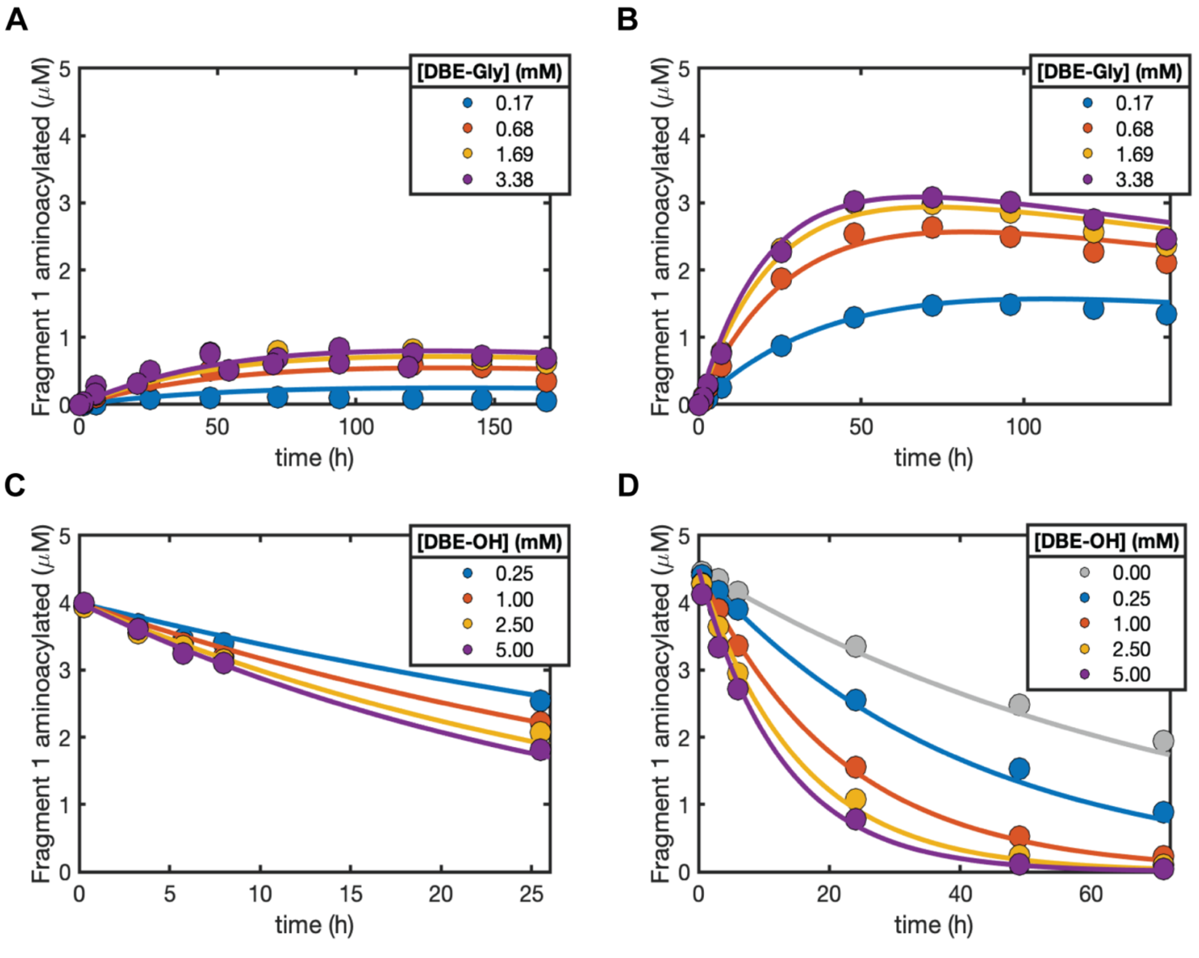


**Fig. S5. Aminoacylation and deacylation activities of the fragmented and the P1 Flexizyme. A:** Aminoacylation activity of the fragments was measured with increasing concentrations of DBE-gly. The reactions contained 5 μM each of the 5′-FAM labeled fragment 1 and 2, and 1 μM of fragment 3. **B:** As in **A**, except the reactions contained 5 μM of the 5′-FAM labeled fragment 1, 5 μM of the model oligonucleotide (Model 2 oligo in Table S1), 0.5 μM fragment 3, and 0.5 μM of the **P1** Flexizyme. **C:** Deacylation activity of the fragments was measured with increasing concentrations of DBE-OH. The reactions were set up as in **A**, except 5 μM of the purified, glycylated and 5′-FAM labeled fragment 1 was used instead of 5 μM of the 5′-FAM labeled fragment 1. **D:** As in **C**, except the reactions contained 5 μM of the glycylated and 5′-FAM labeled fragment 1, 5 μM of the model oligonucleotide (Model 2 oligo in Table S1), 0.5 μM fragment 3, and 0.5 μM of the **P1** Flexizyme. The traces in all four panels corresponded to computationally fitted reaction rates.

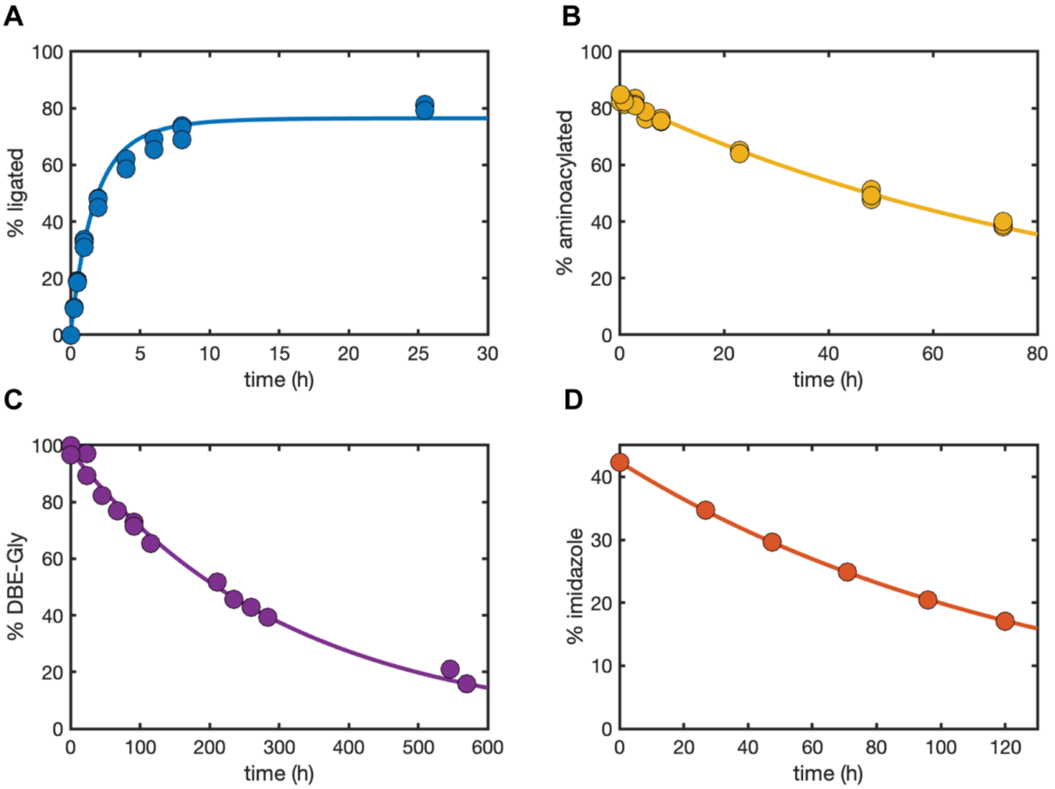


**Fig. S6. Kinetics of the loop-closing ligation and hydrolyses in the autocatalytic reaction network. A:** The % ligated product was measured by PAGE in a reaction containing 5 μM glycylated and 5′-FAM labeled fragment 1, 5 μM 5′-phosphorimidazolide activated fragment 2, and 3.38 mM DBE-gly. The absence of fragment 3 prevents re-aminoacylation of the hydrolyzed fragment 1 and permits the measurement of the isolated ligation rate. **B:** Hydrolysis of the glycylated and 5′-FAM labeled fragment 1 in the presence of 5 μM fragment 2 and 3.38 mM DBE-gly was measured by acidic PAGE. Re-aminoacylation was prevented as in **A**. **C:** Hydrolysis of DBE-gly to DBE-OH in 100 mM imidazole pH 8 and 5 mM $MgCl_2$ was measured by $^1H$ NMR. **D:** Hydrolysis of the 5′-phosphorimidazolide activated model oligonucleotide (Model 2 oligo in Table S1) was measured by analytical HPLC. The traces in all four panels corresponded to computationally fitted reaction rates.

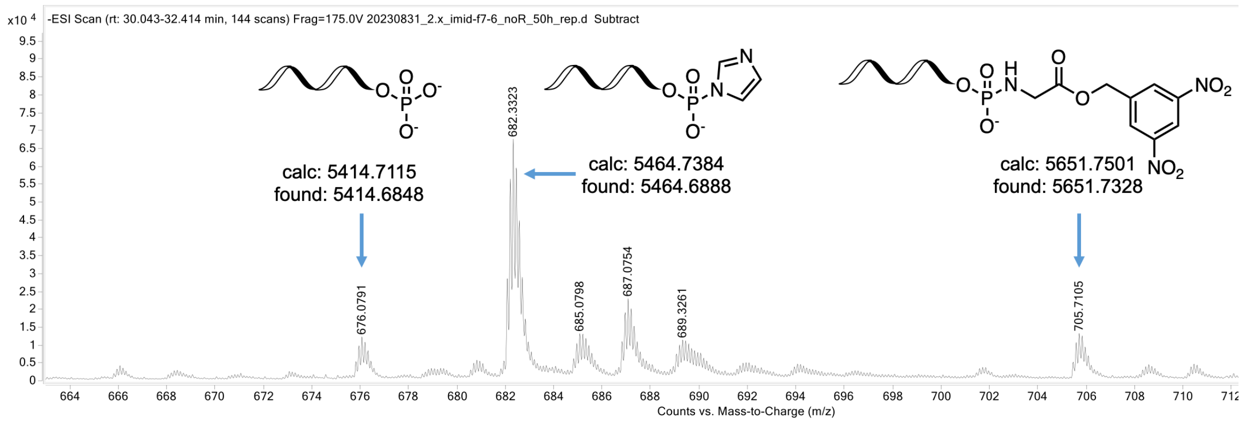


**Fig. S7. TOF analysis of the 5′-phosphoramidate formation in the presence of DBE-gly.** The 5′-phosphorimidazolide activated fragment 2 was incubated with 3.38 mM DBE-gly in 100 mM imidazole pH 8 and 5 mM $MgCl_2$ for 72 hours. A single signal with ion counts above the noise was found in the total ion chromatogram. Shown is the extracted ion chromatogram that contained the indicated fragment 2 derivatives.

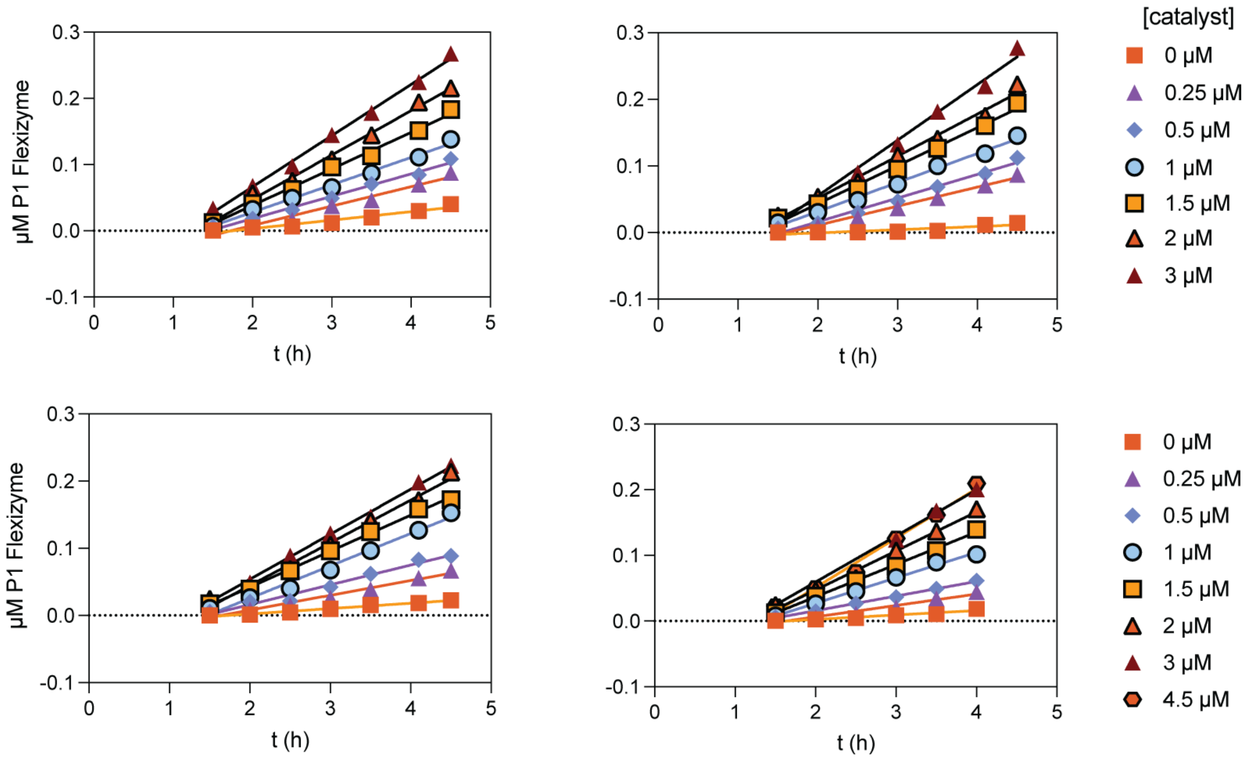


**Fig. S8. Spike-in experiment for determining the reaction order with respect to catalyst.** Four independent experiments measuring the amount of ligated P1 chimeric Flexizyme formed in the initial stages of the autocatalytic assembly reaction with a range of preformed P1 Flexizyme spike-in concentrations. The reaction conditions were identical to those used in Fig. 3. Beyond the 4.5-hour timepoint, the product formation is no longer linear. The initial rates of the reaction in μM $h^{-1}$ were obtained from the slope of the linear fit.

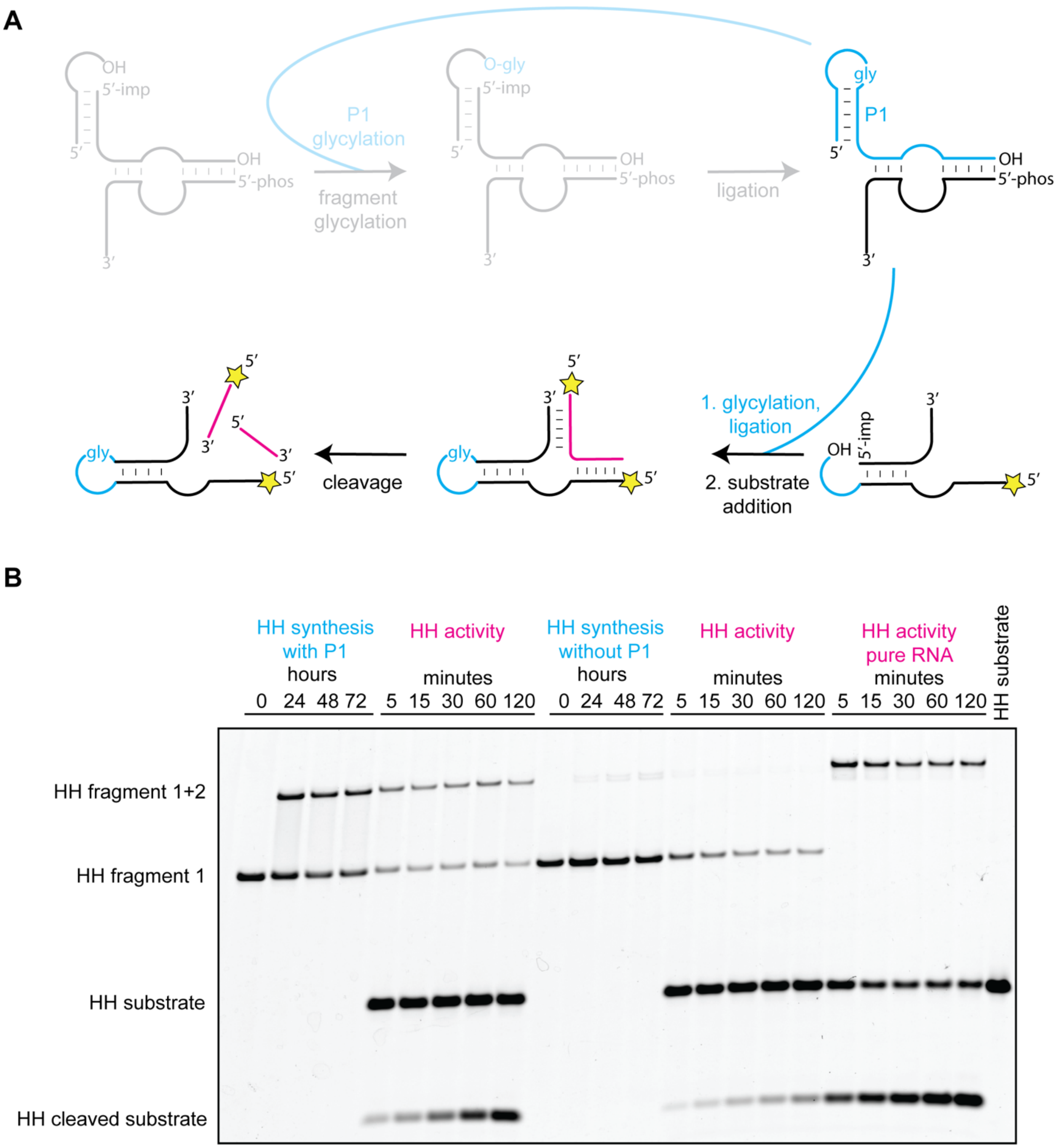


**Fig. S9. Chimeric hammerhead synthesis and activity. A:** The chimeric P1 Flexizyme is first synthesized in the autocatalytic self-assembly cycle (greyed out), then used to aminoacylate the hammerhead fragments, which after ligation form a chimeric stem-loop. The hammerhead substrate is pink. Yellow star represents a 5′-FAM label. Note that the P1 ribozyme was not 5′-FAM labeled. **B:** Denaturing PAGE of the hammerhead assembly and activity. Conditions for HH synthesis for the same as in Fig. 5, except in the case of ‘HH synthesis without P1’ where no P1 ribozyme was added. Conditions for HH activity were the same as in Fig. 5.

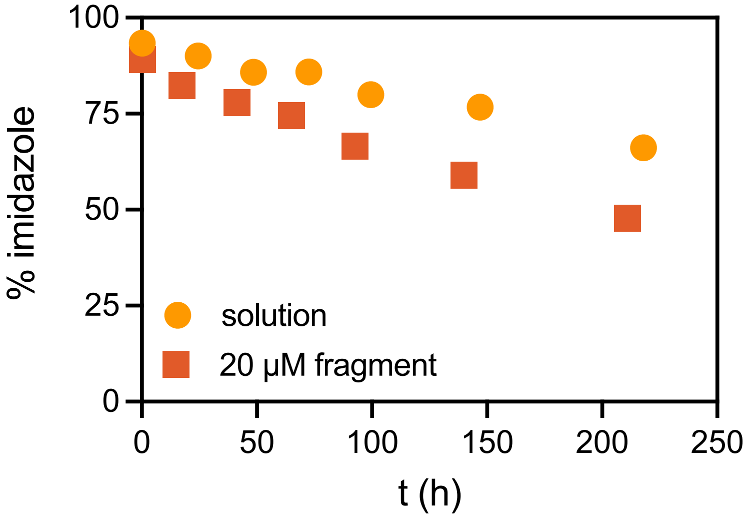


**Fig. S10. Flexizyme-catalyzed 5′-phosphoramidate fragment 2 synthesis.** The solution reaction contained 100 mM imidazole pH 8, 5 mM $MgCl_2$, 3.38 mM DBE-gly, and 20 μM 5′-phosphorimidazolide fragment 2. The 20 μM fragment reaction additionally contained 20 μM of the periodate-treated fragment 1, and 5 μM fragment 3. Due to the chromatographic overlap of the 5′-phosphoramidate and the 5′-phosphate (the hydrolysis product of the 5′-phosphorimidazolide), we monitored the disappearance of the 5′-phosphorimidazolide fragment 2 via analytical HPLC. The 20 μM fragment reaction accelerated the disappearance of the 5′-phosphorimidazolide fragment 2 and we used the difference in the rates between the 20 μM fragment and solution reactions to estimate the rate of 5′-phosphoramidate synthesis.

**Table S1. RNA sequences used in this work**. Imp represents imidazole activated phosphate.

| Name | Sequence |
|---|---|
| Fragment 1 (F1) | 5′-fluorescein-GGACCUGAGAAA |
| Fragment 2 (F2) | 5′-phos/Imp-GGUCCCGCAUCCCAGUC |
| Fragment 3 (F3) | 5′-phos-GACUGGUACAUGGCGUUAUUCUC |
| Fragment 2_overhang | 5′-phos/Imp-GGUCCCGCAUCCCAGUCUGAGAAA |
| Fragment 2_5BP | 5′-phos-GGUCCCGCAUCCCAUCUGAGAAA |
| Fragment 3_5BP | 5′-phos-GAUGGUACAUGGCGUUAUUCUC |
| **P1+P2** | 5′-fluorescein-GGACCUGAGAAA-gly-GGUCCCGCAUCCCAGUCUGAGAAA-gly-GACUGGUACAUGGCGUUAUUCUC |
| **P1** | 5′-fluorescein-GGACCUGAGAAA-gly-GGUCCCGCAUCCCAGUC |
| **P2** | 5′-phos-GGUCCCGCAUCCCAGUCUGAGAAA-gly-GACUGGUACAUGGCGUUAUUCUC |
| Model 2 | 5′-phos-GGUCC |
| dFx_S7 | 5′-fluorescein-GGACCUGAGAAA GGUCCCGCAUCCCAUCUGAGAAA GAUGGUACAUGGCGUUAUUCUC |
| **P1** unlabeled | 5′-OH-GGACCUGAGAAA-gly-GGUCCCGCAUCCCAGUC |
| HH1 | 5′-fluorescein-AAGCACACUGAUGAGCCUUGAGAAA |
| HH2 | 5′-phos/Imp-AGGCGAAACGAU |
| HH substrate | 5′-fluorescein-AGAUCGUCUGUGC |
| HH all-RNA | 5′-fluorescein-AAGCACACUGAUGAGCCUUGAGAAAGGCGAAACGAU |
